# Minimal and mild hearing loss in children: Association with auditory perception, cognition, and communication problems

**DOI:** 10.1101/723635

**Authors:** David R. Moore, Oliver Zobay, Melanie A. Ferguson

## Abstract

**Objectives:** ‘Minimal’ and ‘mild’ hearing loss are the most common but least understood forms of hearing loss in children. Children with better ear hearing level as low as 30 dB HL have a global language impairment and, according to the World Health Organization, a “disabling level of hearing loss”. We examined in a population of 6 - 11 year olds how hearing level ≤ 40.0 dB HL (1 and 4 kHz pure tone average, PTA, threshold) related to auditory perception, cognition and communication.

**Design:** School children (n=1638) were recruited in four centres across the UK. They completed a battery of hearing (audiometry, filter width, temporal envelope, speech-in-noise) and cognitive (IQ, attention, verbal memory, receptive language, reading) tests. Caregivers assessed their children’s communication and listening skills. Children included in this study (702 male; 752 female) had four reliable tone thresholds (1, 4 kHz each ear), and no caregiver reported medical or intellectual disorder. Normal hearing children (n=1124, 77.1%) had all four thresholds and PTA < 15 dB HL. Children with ≥ 15 dB HL for at least one threshold, and PTA < 20 dB (n=245, 16.8%) had Minimal hearing loss. Children with 20 ≤ PTA < 40 dB HL (n=88, 6.0%) had Mild hearing loss. Interaural Asymmetric hearing loss (|Left PTA – Right PTA| ≥ 10 dB) was found in 28.9% of those with Minimal and 39.8% of those with Mild hearing loss.

**Results:** Speech perception in noise, indexed by VCV pseudoword repetition in speech modulated noise, was impaired in children with Minimal and Mild hearing loss, relative to Normal hearing children. Effect size was largest (d=0.63) in Asymmetric Mild hearing loss and smallest (d=0.21) in Symmetric Minimal hearing loss. Spectral (filter width) and temporal (backward masking) perception were impaired in children with both forms of hearing loss, but supra-threshold perception generally related only weakly to PTA. Speech-in-noise (nonsense syllables) and language (pseudoword repetition) were also impaired in both forms of hearing loss and correlated more strongly with PTA. Children with Mild hearing loss were additionally impaired in working memory (digit span) and reading, and generally performed more poorly than those with Minimal loss. Asymmetric hearing loss produced as much impairment overall on both auditory and cognitive tasks as Symmetric hearing loss. Nonverbal IQ, attention and caregiver-rated listening and communication were not significantly impaired in children with hearing loss. Modelling suggested that 15 dB HL is objectively an appropriate lower audibility limit for diagnosis of hearing loss.

**Conclusions:** Hearing loss between 15 - 30 dB PTA is, at ~20%, much more prevalent in 6-11 y.o. children than most current estimates. Key aspects of auditory and cognitive skills are impaired in both symmetric and asymmetric minimal and mild hearing loss. Hearing loss < 30 dB HL is most closely related to speech perception in noise, and to cognitive abilities underpinning language and reading. The results suggest wider use of speech-in-noise measures to diagnose and assess management of hearing loss and reduction of the clinical hearing loss threshold for children to 15 dB HL.

## INTRODUCTION

### Levels and prevalence of hearing loss in children

Disabling hearing loss in children is currently defined by the World Health Organization (2012) as pure tone average threshold, PTA > 30 dB hearing level (HL) in the better hearing ear, occurring worldwide in 1.7% of 0-14 year olds (y.o.). However, lower levels of hearing loss (PTA 15-30 dB HL), both unilateral and bilateral, have been associated with educational and communicative difficulties (Bess et al. 1998), and the prevalence of this lower level hearing loss is high. In one prominent study (Niskar et al. 1998), 14.9% of 6-19 y.o. in a large US-based population (NHANES) had a hearing loss > 15 dB HL in at least one ear. Most of this hearing loss was unilateral, ‘slight’ (PTA 16-25dB HL) and higher frequency (3-8 kHz). Other studies broadly concur with these findings. For example, Bess and colleagues (1998) found 137/1218 (11.3%) of a sample of school children (8-15 y.o.) had hearing loss (PTA 15-40 dB HL), of whom about half (n=66) had sensorineural loss and half had conductive or other losses. Of those with sensorineural loss, 37 (56%) had a ‘unilateral’ loss, whereas the remainder had a ‘mild bilateral’ or ‘high frequency’ hearing loss. In a recent meta-analysis, unilateral, mild bilateral, and high frequency loss (20-40 dB HL) were again identified as the most common configurations of lower level sensorineural hearing loss in children (Bess et al. 1998; Winiger et al. 2016).

More generally, in both adults and children, there appears to be no objective rationale for a lower limit to what is called hearing loss, with a range of descriptors for different hearing threshold levels and frequencies (Timmer et al. 2015). The same is true for the terms ‘minimal’ and, to a lesser extent, ‘mild’ hearing loss, with no consensus on definitions. It seems certain that any specific limit will be a simplification in any case, since loss of auditory function (hearing impairment) is individually and, across a population, continuously variable, and includes impairments that are not well predicted by pure tone detection threshold (Pienkowski, 2017; Le Prell, 2019). Prevalence estimates for hearing loss ≤ 40 dB HL in children vary widely from one study to another and depend on sampling strategy, type and level of hearing loss, and demographic factors (Niskar et al. 1998; Wake et al. 2006; WHO, 2012; Winiger et al. 2016). Nevertheless, it is clear that many more children have a hearing loss in the range 15-40 dB HL than those who have a more severe form of hearing loss (Feder et al. 2017).

### Consequences of hearing loss

The consequences of lower level hearing loss for the development of language and other key life skills in children have been examined in several recent large-scale studies. These studies have focussed on cases of hearing loss detected from neonatal hearing screening (Moeller & Tomblin, 2015b; Ching et al. 2017). While children with hearing loss from 20-40 dB HL made up 46% of those with permanent hearing loss in one study (Fitzpatrick et al. 2014), neonatal screening generally does not detect hearing loss less than about 30 dB HL (Norton et al. 2000). However, a ‘global language score’ well under the mean for normally-hearing children was found for 5 year olds with a better-ear hearing level of 30 dB (Ching et al. 2017). Together, these data suggest a likely high prevalence and significant impact of hearing loss < 40 dB HL.

There are limited data on the consequences of hearing loss < 30 dB HL for auditory perception, listening, cognition, speech and communication in children, mainly due to insensitive screening, low referral, and low elective attendance of affected individuals at clinics. Existing population data are often restricted to comparisons between PTA and demographic metrics (e.g. age, sex, laterality; A.J. Hall et al. 2011). One large study (Wake et al. 2006) found that bilateral loss of 16-40 dB HL was *not* associated with impaired language, reading, behaviour or health-related quality of life, although phonologic short-term memory was reduced. That study stands in contrast to most others suggesting lower level hearing loss can affect broader abilities. As above, Bess and colleagues (1998) showed early language and literacy difficulties and persistent problems with communication and social skills among a more heterogeneous sample of children. Laboratory studies of children diagnosed with ‘mild/moderate’ hearing loss (mostly PTA > 35 dB HL) have also shown associations with poor auditory (Hall et al. 2012), language (Briscoe et al. 2001) and comprehension (Lewis et al. 2015) skills. Overall, despite the prevalence and importance of hearing loss between 15-40 dB HL, and the many studies published, most of the available data are from the upper end of this range (30-40 dB HL). It is also unclear what the minimum hearing loss should be, in one or both ears, to trigger intervention (McKay et al. 2008).

### Asymmetric hearing loss

Unilateral hearing loss, previously considered a form of minimal hearing loss but not a major clinical concern (Bess et al., 1998), has recently received more attention. That attention has mainly focused on single sided deafness, a severe or profound and important form of unilateral hearing loss, but not considered further here (see Anne et al. 2017; van Wieringen et al. 2019, for recent reviews). Some studies have also included limited data on mild unilateral hearing loss, but those studies have typically recruited through neonatal screening and, consequently, the samples have been very small (e.g. Fitzpatrick et al. 2019). Since there is a continuum of hearing sensitivity in either ear of children with PTA ≤ 40 dB HL, we used the term ‘Asymmetric’ hearing loss in this study.

### Study design considerations

As convenient working labels, we adopted the terms ‘Minimal’ hearing loss, used previously to describe a variety of hearing loss including PTA of 15 - 20 dB HL in either ear, ‘Mild’ hearing loss, often PTA of 20 – 40 dB HL (BSA, 2011), and ‘Normal’ hearing, sometimes used to describe those with thresholds in both ears < 15 dB HL, but more usually defined as a higher threshold level. ‘Asymmetric’ loss has most frequently been used to designate interaural PTA differences ≥ 15 dB, although ≥ 10 dB and ≥ 20 dB have also been used (Durakovic et al. 2018). In this study, we adopt relatively low level but specific criteria for these labels to include all cases that may be of concern. However, because of the continuous nature of these measures, we used continuous analysis in some instances.

Critical to success of this study was the adoption of a rapid means for evaluating hearing level that was adequately accurate and usable in low-noise environments (a ‘quiet room’ within a school) rather than inside a sound booth. We chose a method of automated (‘Békésy’) audiometry to establish thresholds for 1 and 4 kHz tones only, presented monaurally through sound attenuating headphones. Considerable pre-experimental piloting in and out of a sound booth and post-recording analysis was performed to establish the reliability of this technique and its ability to measure thresholds accurately (± 5 dB) down to 10 dB HL. A key requirement was that the participant needed to signal at least 6 reversals of roving level for each threshold to be recorded and computed. Comparison with published reports (Beahan et al. 2012) using conventionally recorded audiometric data in children over an age range including that used here showed a comparable level of test-retest reliability (~ 5 dB), and its modest (~ 2 dB) improvement with age between 6 – 11 y.o.

Audiometric thresholds are not necessarily a good predictor of speech perception, particularly in acoustically challenging environments (Pienkowski 2017; Le Prell 2019). This may be due to the unique auditory requirements of processing speech and to individual variation in the additional cognitive load of recognizing speech-in-noise, compared with the detection of tones-in-quiet. Studies in adults clearly show that speech perception is partly influenced by cognitive factors, separate from audiometric sensitivity (Füllgrabe et al. 2015; Heinrich et al. 2015), and that those with stronger cognitive skills tend to have better speech reception thresholds than those who have poorer cognitive skills (Moore et al. 2014). However, we do not currently know fully to what extent children with minimal and mild hearing loss have a broader range of functional deficits, how those deficits relate to their hearing level, whether the children sometimes ‘compensate’ for their hearing loss, or at what level or asymmetry of loss other difficulties occur.

We conducted a large, multi-center population study of hearing, perceptual, cognitive, and communicative and listening abilities in primary (elementary) schools across the UK. This study was aimed initially at children who were audiometrically ‘normal’ (all tone thresholds ≤ 25 dB HL; (Moore et al. 2010). However, a significant, additional number of children who had one or more thresholds above this level also completed the full range of tests. For the study reported here, we recalculated pure tone thresholds of all children in the study using a stricter and more objective set of criteria. Most of the children (77%) had Normal hearing and a mere 1.7% had PTA ≥ 30 dB HL. In this respect, the study population was almost a mirror image of those discussed above. For those with hearing loss, data are additionally reported for Symmetric and Asymmetric loss. The specific aims of the study were to (i) relate hearing level across a broader range of PTA (< 0 to ≤ 40.0 dB HL) to auditory perception, cognition, communication and listening, (ii) make an objective determination of the minimum HL at which difficulties occur, (iii) examine functional ‘compensation’ for hearing loss, testing the hypothesis that children with stronger cognitive skills tend to be less impaired by their hearing loss than others, and (iv) compare the effects of symmetric and asymmetric hearing loss.

## METHODS

### Design and Population

The study was stratified by age (6:0-11:11, years:months), sex and socioeconomic status to reflect UK demographics. Our aim was to perform a wide-ranging evaluation of skills in a large population within a school setting and to acquire complete test data on an individual child within one hour. Extensive, preliminary, lab-based trialling suggested this was possible (Moore et al. 2011).

All participants used English as the main home language. Written consent and background questionnaire results (audiology, education, medical history) were obtained from caregivers following invitation packs mailed to 8044 homes. Ethical and associated approvals were received from host National Health Service Trusts and local educational authorities.

Of 1638 children from 44 mainstream primary schools in Cardiff, Exeter, Glasgow and Nottingham participating in a 1-hour test session, data from 1457 (752 female) children had four reliable tone thresholds (2 frequencies, 1 kHz and 4 kHz x 2 ears), and no stated medical or intellectual disorder as determined by a background questionnaire. Normal hearing children (n=1124, 77.1%) had all thresholds < 15 dB HL at 1 kHz and 4 kHz in both ears. Children with ≥ 15 dB HL for at least one threshold, and < 20 dB PTA (n=245, 16.8%) were designated Minimal loss^1^, and those with 20 ≤ PTA < 40 dB PTA (n=88, 6.0%) Mild loss. Interaural, asymmetric hearing loss (|Left PTA – Right PTA| ≥ 10 dB) was found for 71/245 (28.9%) of those with Minimal and 35/88 (39.8%) of those with Mild loss. Among Normal hearing children, 27/1124 (2.4%) had asymmetric hearing ≥ 10 dB.

### Data Collection

Full details of all measures used in this study have been published (Moore et al. 2010). Auditory perception tasks were developed, piloted and extensively tested as documented in other studies from our group, and most other methods used here were similarly used in those studies (Ferguson et al. 2011; Moore et al. 2011). Children were tested by university graduates who received extensive training on the specific requirements of this study. Testing took place in quiet locations in the children’s schools using PC laptop computers running IHR-STAR ‘IMAP’ task control and signal generation software (Barry et al. 2010). Test sessions included modules to measure pure-tone thresholds (Fig. 1), auditory perception (Fig. 2) and cognition. Caregivers completed the Children’s Communication Checklist, 2^nd^ Edition (CCC-2; Bishop 2003, a well-established and validated instrument to assess speech and language skills, the Children’s Auditory Processing Performance Scale (CHAPPS; Smoski 1992), a teacher or parent report of listening skills commonly used in studies of auditory processing disorder (APD), and a background questionnaire devised for the study asking questions about the child’s auditory (e.g. Has the child received grommets? ‘PE tubes’), neurological and educational history. Both the CCC-2 and the CHAPPS were mailed to caregivers and returned in stamped, return addressed envelopes.

**Fig. 1:**
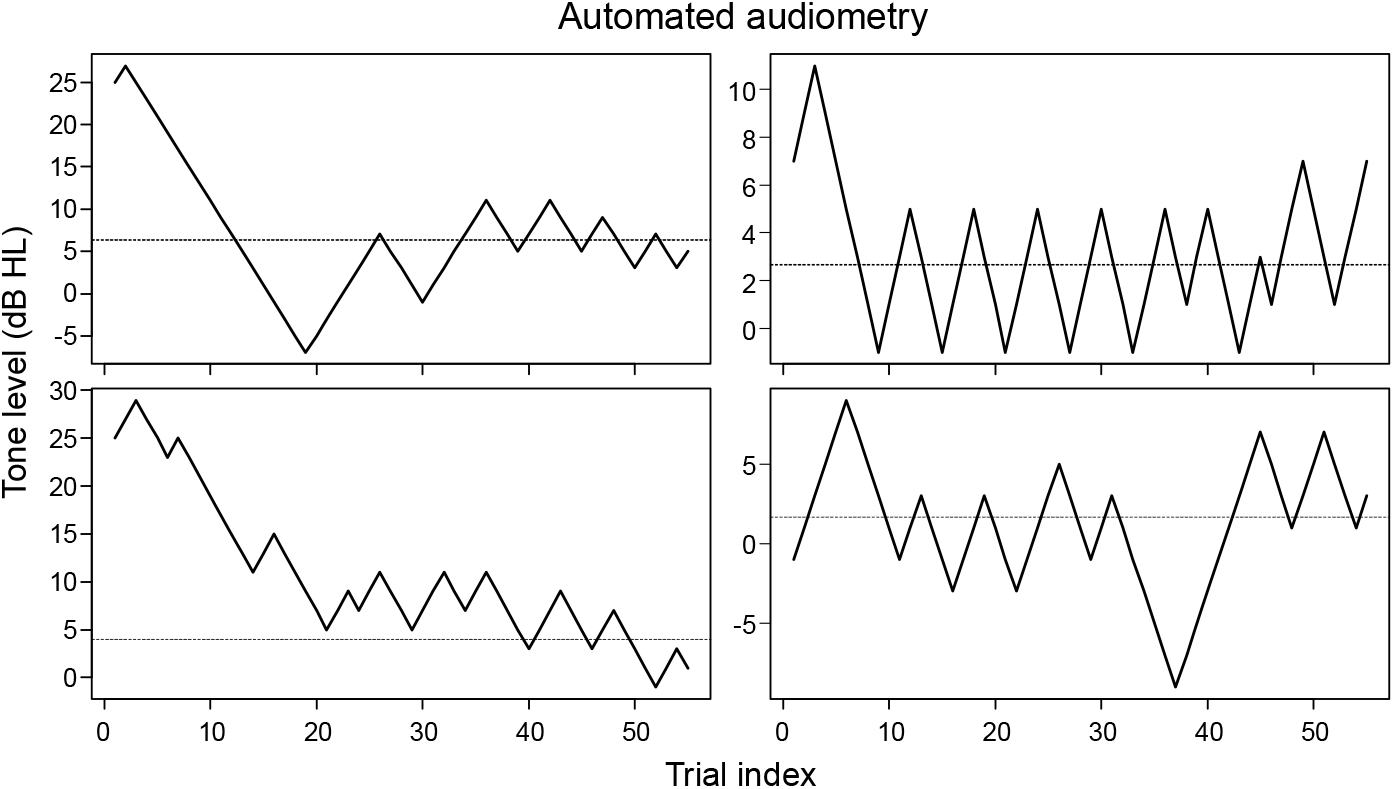
Testing pure tone sensitivity. Examples of automated (Békésy) audiometry showing tone level changes across trials resulting in ‘reversals’ of the direction of level change contingent on the child signalling loss or gain of audibility. To be included in the analysis sample, each child had to make at least 6 such reversals for each of four tone/ear combinations. The dotted line shows the derived pure tone threshold.

**Fig. 2:**
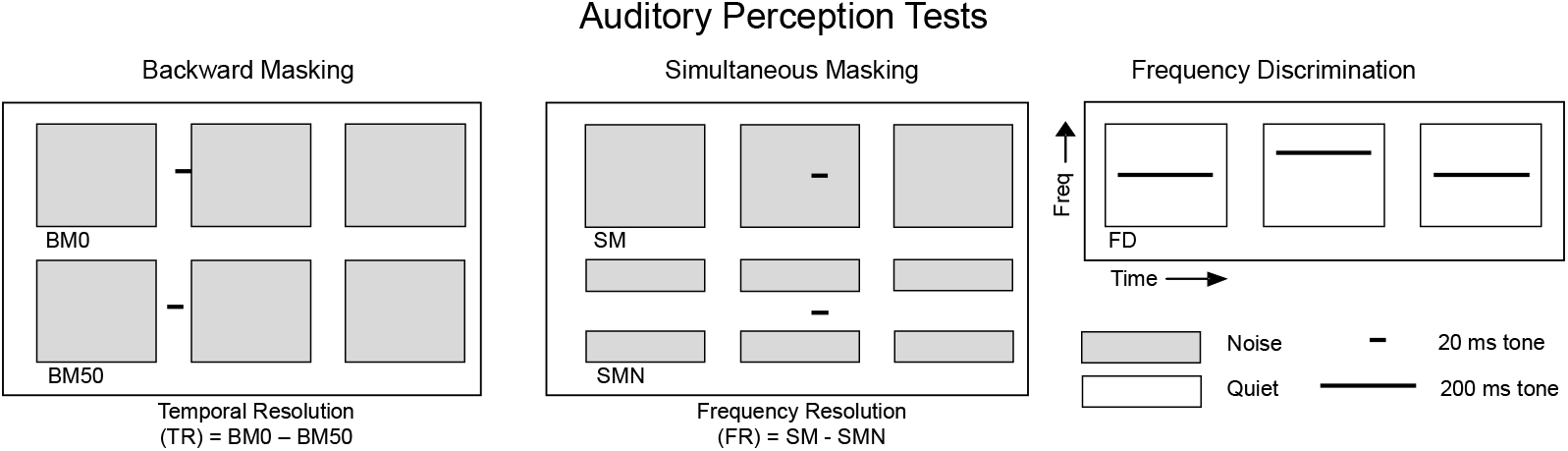
Auditory perception (AP) was measured psychoacoustically. Schematic of individual (BM0, BM50, SM, SMN, FD) and derived (TR, FR) AP tests. Boxes designate sound presentation intervals of noise or quiet. Lines are target tones. For each test, an individual ‘trial’ involved presentation of three successive intervals (each 300 ms) defined either by the presence of a band-limited noise (grey; Backward and Simultaneous Masking) or a 200 ms tone (Frequency Discrimination, FD). For the masking tests, a short (20 ms) 1 kHz tone was presented before or during the noise. For the BM tests, the tone was presented immediately before (BM0) or with a delay of 50 ms before (BM50) the noise. For the SMN test the noise contained a spectral notch centred on the tone frequency (freq). For the FD test one of the three tones was presented at a higher frequency than the other two, identical tones. For all tests the interval containing the odd stimulus (shown in the 2^nd^ interval here) varied randomly from one trial to the next. The listener’s task was to indicate with a button press which interval contained the odd stimulus. Tone level (masking tests) or relative frequency (FD test) varied adaptively with the listener’s performance. Each test took about 5 minutes to complete 40 trials with a short break after the first 20 trials.

### Tone Thresholds – Audiometry

An automated audiometer (software supplied by Dr M. Lutman, ISVR, University of Southampton), running under IHR-STAR ‘IMAP’, was used to present 1 and 4 kHz tones monaurally to each ear via Sennheiser supra-aural, closed-back HD 25-1 headphones. Each pair of headphones and their host computer was checked for distortion and calibrated for sound pressure level at the MRC Institute of Hearing Research (IHR) according to ISO-389.1 (2007). Children were asked to press a button whenever they heard a ‘beeping’ sound, to keep the button pressed as long as they could hear the sound, and to stop pressing the button as soon as they could no longer hear the sound. Threshold was the mean of the last 6 reversals, and only data from children who completed 6 or more reversals for both frequencies and both ears were included in this study. A few examples are shown in Fig. 1.

### Auditory Perception

Four of the auditory perception (AP) tests involved detection of a 20 ms 1 kHz tone pulse (10 ms cos^2^ ramps; initially 75 or 90 dB SPL), adaptively varied (3-down, 1-up) in level from an initial 90 dB SPL, presented to both ears before or simultaneously with a fixed spectrum level (30 dB/Hz), 300 ms band-pass (600 – 1400 Hz) noise masker (Fig. 2). Children were asked to identify (label) and remember, until responding at the end, which of three successively presented sounds was the ‘odd-one-out’ (i.e. contained the tone). Response was via three large, coloured buttons corresponding to the three sounds. Two ‘individual’ versions of a test measured tone detection thresholds in a backward masking procedure; the tone was presented either immediately (BM0) or 50 ms (BM50) before the noise. Two other individual test versions measured tone thresholds presented in the same masking noise simultaneously, either with (SMN) or without (SM) a spectral notch (400 Hz) centred on the frequency of the tone (1 kHz). By subtracting thresholds from each version of the same test, we produced ‘derived’ measures of temporal (TR) and frequency resolution (FR), respectively, each based on two data points. If we assume the procedural demands (attention, working memory, executive function) of each version of the tests were the same, this subtraction process should cancel out the contribution of those demands, leaving relatively pure measures of sensory function (Moore, 2012; Dillon, 2014). The fifth AP test, frequency discrimination (FD), presented three successive 200 ms tones diotically on each trial. Here, the odd-one-out had a higher pitch varied adaptively with the standards fixed at 1 kHz. A speech-in-noise test consisted of vowel-consonant-vowel (VCV) pseudowords in speech modulated noise (Ferguson et al. 2011). Pseudowords were made by placing one of three vowels ([a:], [i:], [u:]) either side of one of 20 consonants, yielding 60 possible combinations (e.g. “iji” and “unu”). Each pseudoword test list contained 20 items, and scoring was based on correct repetition of the consonant. Noise was fabricated from a single male talker (ICRA-5; Dreschler et al 2001), presented at 65 dBA. Children repeated verbally the VCV target as its level varied adaptively around that of the constant-level noise. Speech reception thresholds (SRT) were the mean reversal level of the VCV in dBA.

### Cognition

Cognitive tests were age-appropriate, standardized measures of aspects of developmental ability and learning that are often-reported outcome measures in studies of communication disability - nonverbal reasoning (NVIQ; Matrices Reasoning; Wechsler 1999), working memory (digit span; (Wechsler 1991), phonological processing and verbal working memory (‘language’; pseudoword repetition, NEPSY; Korkman 1998), and reading accuracy and fluency (TOWRE words and pseudowords; Torgeson et al. 1999). Attention was assessed through performance variability in the AP tests (‘intrinsic’ attention) and through ‘extrinsic’ tests of auditory and visual ‘phasic alertness’, the IHR Cued Attention Test (IHR-CAT). These attention measures were developed specifically for the studies presented here and in Moore et al. (2010), where they are described and discussed extensively.

### Communication and Listening

The CCC-2 has 70 items arranged in 10 scales (Bishop, 2003). We used the General Communication Composite (GCC) measure that summed scores on 8 scales covering language skills and nonverbal communication. The CHAPPS has a single composite score for 36 items covering listening in different environments, auditory memory, and auditory attention.

### Analysis

Outcome measures were compared between normal hearing and hearing loss categories using general linear models (ANOVA, ANCOVA), controlling for age where appropriate (i.e. AP measures, speech-in-noise, attention). For post-hoc comparisons, Fisher’s LSD test was used (Maxwell & Delaney, 2004). Effect sizes were measured by Cohen’s d, with age considered as extrinsic and cognition as intrinsic covariates (Maxwell & Delaney, 2004). Logistic regression was used to investigate the relation between hearing loss on poor test performance, defined as < 5% of mean Normal hearing performance. Independence of factors in two-way contingency tables was tested with chi-square tests. P-values below 0.05 were considered statistically significant. No corrections were made for multiple comparisons. Further details of analysis are provided below as Supplementary Information.

## RESULTS

Children with Minimal and Mild, Symmetric and Asymmetric hearing loss (Table 1) had significantly reduced performance relative to Normal hearing children on speech-in-noise hearing, some spectral and temporal auditory perception, and cognitive skills, but not on caregiver evaluations of communication or listening (Table 2).

**TABLE 1.** Hearing loss and pure tone thresholds across age. Mean (s.d.) hearing level at each frequency, interaural symmetry, and number and proportion of children in each hearing category from 6 – 11 years. Categories were Symmetric and Asymmetric, Minimal and Mild hearing loss, and Normal hearing. 1kHz and 4 kHz mean thresholds (dB HL, averaged across ears) for all children at each age are shown at bottom of table.

| Category/Age | Mean HL<br>dB (s.d.) |  | Mean Asym<br>dB (s.d.) | 6y.o. | 7y.o. | 8y.o. | 9y.o. | 10y.o. | 11y.o. | Total |
| --- | --- | --- | --- | --- | --- | --- | --- | --- | --- | --- |
|  | 1 kHz | 4 kHz |  |  |  |  |  |  |  |  |
| Min Symm | 17.6(4.0) | 15.3(2.0) | 4.7 | n=26 | 41 | 30 | 26 | 34 | 17 | 174 |
| Min Asymm | 16.2(4.9) | 14.2(3.2) | 15.5 | 16 | 11 | 10 | 15 | 8 | 11 | 71 |
| Mild Symm | 26.4(5.6) | 25.7(4.4) | 4.7 | 14 | 12 | 12 | 6 | 8 | 1 | 53 |
| Mild Asymm | 26.5(6.2) | 26.0(4.5) | 22.1 | 9 | 3 | 6 | 5 | 8 | 4 | 35 |
| Norm hearing | 8.3(4.7) | 6.1(3.8) | 3.5 | 148 | 203 | 185 | 231 | 224 | 133 | 1124 |
| Total children |  |  |  | 213 | 270 | 243 | 283 | 282 | 166 | 1457 |
| % Minimal |  |  |  | 19.7% | 19.3 | 16.5 | 14.5 | 14.9 | 16.9 | 16.8 |
| % Mild |  |  |  | 10.8 | 5.6 | 7.4 | 3.9 | 5.7 | 3.0 | 6.0 |
| % Normal |  |  |  | 69.5 | 75.2 | 76.1 | 81.6 | 79.4 | 80.1 | 77.1 |
| 1 kHz mean threshold |  |  |  | 11.9 | 10.6 | 10.9 | 10.1 | 10.0 | 9.3 |  |
| 4 kHz mean threshold |  |  |  | 8.6 | 6.2 | 6.0 | 5.0 | 5.8 | 5.3 |  |

**TABLE 2.** Statistical comparisons between children with and without hearing loss. Results of overall hearing category comparison, controlling for age (ANCOVA; p), with post-hoc, pair-wise comparisons (p) and effect sizes (Cohen’s d). Symm and Asymm are relevant only to those with minimal or mild hearing loss. Significant overall comparisons (p < 0.05) in bold.

| <u>Minimal hearing loss</u> | Overall<br>p | Symm/Normal<br>d p |  | Asym/Normal<br>d p |  | Symm/Asym<br>d p |  |
| --- | --- | --- | --- | --- | --- | --- | --- |
| **Speech-in-noise (VCV)** | **0.003** | **0.21** | **0.010** | **0.31** | **0.012** | **-0.10** | **0.487** |
| Temporal (BM0) | 0.226 | 0.09 | 0.267 | 0.18 | 0.157 | -0.08 | 0.559 |
| **Temporal (BM50)** | **0.016** | **0.15** | **0.077** | **0.30** | **0.015** | **-0.15** | **0.289** |
| Temporal (TR) | 0.532 | -0.08 | 0.333 | -0.08 | 0.514 | 0.00 | 0.998 |
| Temporal (FD) | 0.574 | 0.05 | 0.528 | 0.12 | 0.371 | -0.06 | 0.673 |
| **Spectral (SM)** | **0.006** | **0.13** | **0.109** | **0.36** | **0.003** | **-0.23** | **0.113** |
| Spectral (SMN) | 0.243 | -0.04 | 0.642 | 0.19 | 0.117 | -0.23 | 0.104 |
| Spectral (FR) | 0.170 | 0.15 | 0.070 | 0.08 | 0.499 | 0.07 | 0.631 |
| Intrinsic Attention | 0.752 | 0.06 | 0.476 | 0.04 | 0.759 | 0.02 | 0.882 |
| Extrinsic Attention | 0.318 | -0.09 | 0.278 | -0.14 | 0.250 | 0.05 | 0.710 |
| Extrinsic Visual | 0.558 | -0.02 | 0.826 | -0.13 | 0.283 | 0.11 | 0.420 |
| Intelligence (NVIQ) | 0.366 | -0.08 | 0.357 | -0.14 | 0.248 | 0.07 | 0.635 |
| Working memory (Digits) | 0.187 | -0.09 | 0.284 | -0.19 | 0.115 | 0.11 | 0.454 |
| **Language (pseudoword rep)** | **0.015** | **-0.24** | **0.004** | **-0.04** | **0.730** | **-0.19** | **0.168** |
| Reading (words) | 0.330 | -0.09 | 0.263 | -0.13 | 0.282 | 0.04 | 0.771 |
| Reading (pseudowords) | 0.241 | -0.12 | 0.144 | -0.12 | 0.333 | 0.002 | 0.987 |
| Communication (GCC)* | 0.519 | -0.08 | 0.378 | -0.11 | 0.422 | 0.03 | 0.853 |
| Listening (CHAPPS)* | 0.589 | -0.09 | 0.366 | -0.08 | 0.569 | -0.005 | 0.977 |

| <u>Mild hearing loss</u> | Overall<br>p | Symm/Normal<br>d p |  | Asym/Normal<br>d p |  | Symm/Asym<br>d p |  |
| --- | --- | --- | --- | --- | --- | --- | --- |
| **Speech-in-noise (VCV)** | **<0.001** | **0.48** | **<0.001** | **0.63** | **<0.001** | **-0.15** | **0.492** |
| **Temporal (BM0)** | **0.008** | **0.24** | **0.101** | **0.48** | **0.006** | **-0.24** | **0.281** |
| **Temporal (BM50)** | **<0.001** | **0.60** | **<0.001** | **0.64** | **<0.001** | **-0.04** | **0.855** |
| Temporal (TR) | 0.291 | -0.22 | 0.122 | -0.06 | 0.731 | -0.16 | 0.475 |
| Temporal (FD) | 0.254 | 0.22 | 0.133 | 0.14 | 0.452 | 0.08 | 0.721 |
| Spectral (SM) | 0.850 | -0.06 | 0.687 | 0.07 | 0.699 | -0.13 | 0.573 |
| **Spectral (SMN)** | **0.003** | **0.36** | **0.011** | **0.43** | **0.020** | **-0.06** | **0.777** |
| **Spectral (FR)** | **0.007** | **-0.26** | **0.071** | **-0.48** | **0.008** | **0.22** | **0.331** |
| Intrinsic Attention | 0.511 | -0.006 | 0.966 | 0.20 | 0.248 | -0.20 | 0.349 |
| Extrinsic Attention | 0.216 | 0.18 | 0.213 | 0.22 | 0.200 | -0.04 | 0.841 |
| Extrinsic Visual | 0.928 | 0.03 | 0.816 | 0.05 | 0.751 | -0.02 | 0.921 |
| Intelligence (NVIQ) | 0.080 | -0.26 | 0.068 | -0.24 | 0.168 | -0.02 | 0.927 |
| **Working memory (Digits)** | **0.002** | **-0.32** | **0.023** | **-0.49** | **0.004** | **0.17** | **0.435** |
| **Language (pseudoword rep)** | **0.004** | **-0.19** | **0.167** | **-0.53** | **0.002** | **0.33** | **0.129** |
| **Reading (words)** | **<0.001** | **-0.47** | **<0.001** | **-0.35** | **0.049** | **-0.12** | **0.577** |
| **Read (pseudowords)** | **0.001** | **-0.48** | **<0.001** | **-0.27** | **0.131** | **-0.21** | **0.346** |
| Communication (GCC)* | 0.438 | -0.16 | 0.336 | -0.19 | 0.375 | 0.02 | 0.927 |
| Listening (CHAPPS)* | 0.125 | -0.30 | 0.088 | -0.25 | 0.241 | -0.05 | 0.860 |

### Hearing Level

The overall distribution of PTA shows two-thirds of thresholds < 10 dB HL, but also 35% ≥ 10 dB HL (Fig. 3). Only a very small proportion of children had PTA ≥ 30 dB HL (n = 17; 1.2%). Younger children (6-8 y.o.) had an increased prevalence (chi-square = 9.02, d.f. = 1; p = 0.003) of higher thresholds at both 1 and 4 kHz than older children (Table 1). This age effect was particularly marked for Mild hearing loss and for 6 y.o. compared with older children. Prevalence of Symmetric and Asymmetric hearing loss did not vary statistically with age (p = 0.19) or hearing loss category (p = 0.06). Mean asymmetry was less for the Minimal Asymmetric hearing loss group (15.5 dB) than for the Mild Asymmetric hearing loss group (22.1 dB). Asymmetry for the Normal hearing group (3.5 dB) did not differ significantly from that of the Minimal and Mild Symmetric groups.

**Fig. 3:**
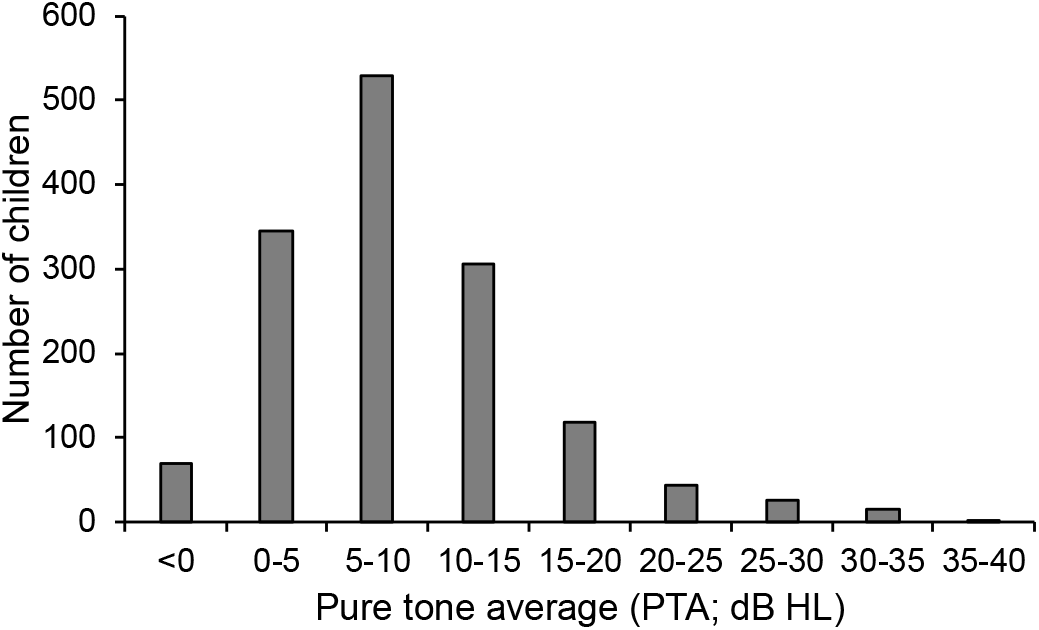
Most children in the sample had low, sensitive pure tone thresholds. Distribution of pure tone average across all 4 threshold measures (2 frequencies x 2 ears) in the whole sample (N = 1457). For each range of PTA, only the lower measure is inclusive (e.g. 0-5 = 0-4.99).

### Auditory Perception (AP)

Mean, age-corrected performance of children on AP tests showed that those with Mild hearing loss generally performed more poorly than those with Minimal loss, as expected (Fig. 4, Table 2). No significant differences were found between Symmetric and Asymmetric hearing loss on any test. Effect sizes for significant Minimal hearing loss were small (d = 0.24 – 0.36) and those for Mild loss ranged from small to moderate, relative to Normal hearing children (d = 0.36 – 0.64; Table 2). For speech-in-noise hearing (VCV), children with both Minimal and Mild hearing loss had significantly higher speech reception thresholds than Normal hearing children. VCV effect sizes were, overall, the largest for any test. Children with Mild, Asymmetric hearing loss had the largest deficits for VCV and for individual spectro-temporal measures (BM0, BM50, SMN), and the only significant impairment on a derived measure (Frequency Resolution; see Methods for explanation).

**Fig. 4:**
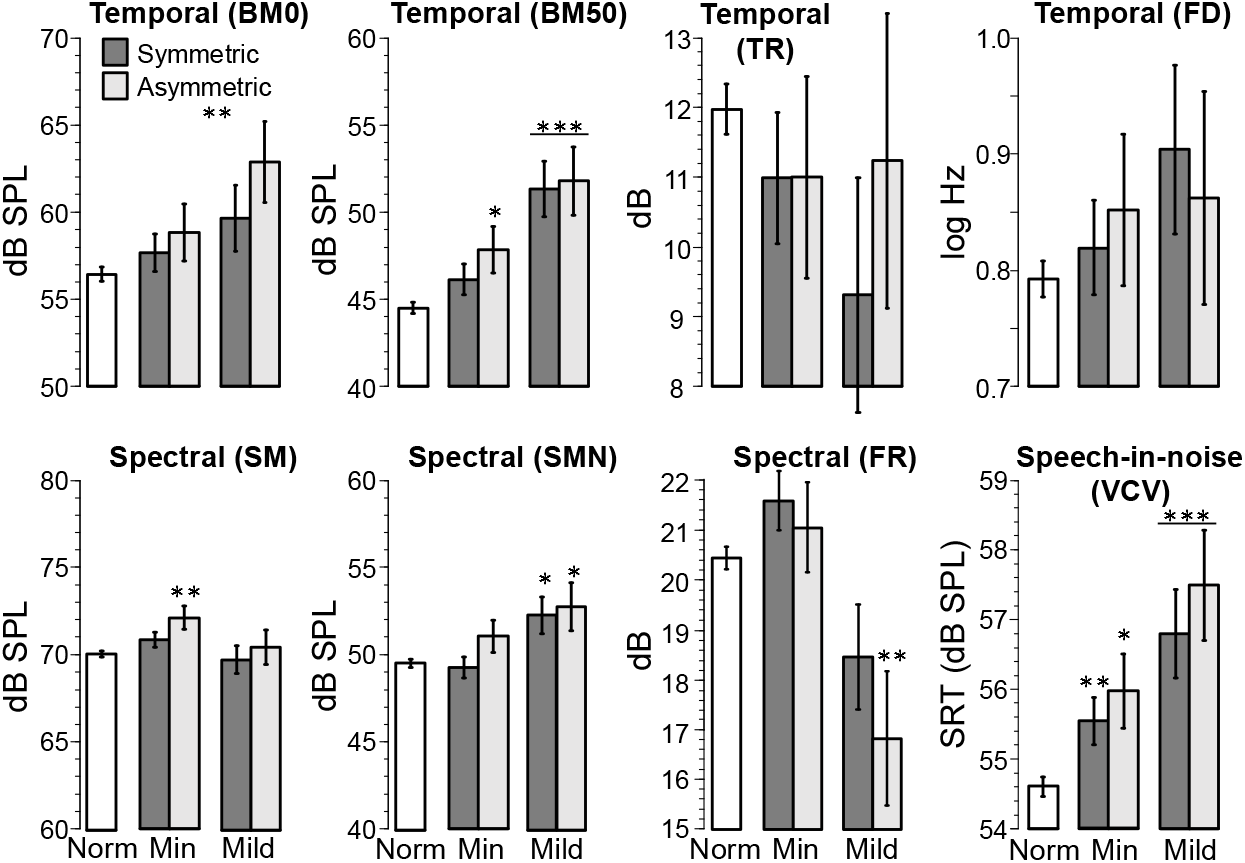
Auditory perception was reduced in children with hearing loss. Histograms show mean (± s.e.m.), age-corrected threshold for AP tests (Fig.2). Age-corrections were estimated from ANCOVA models as expected scores at the overall mean age. Note that for most tests (BM0, BM50, FD, SM, SMN, VCV) a smaller value indicates better performance, but for TR and FR, a larger value indicates better performance. Stars indicate statistically significant differences between hearing loss and Normal hearing categories. * p < 0.05, ** p < 0.01, *** p < 0.001 (See Table 2 for details).

We found only weak correlations between AP measures and four threshold PTA across the whole sample (r = 0.034–0.199; Table 3). These weak correlations could reflect the high proportion of normally hearing children in the population. They could also suggest that a third factor, for example ability to perform AP tests, was affected by hearing loss. Alternately, poorer performance on the pure tone test may have resulted in disproportionate allocation of general poor performers to the Symmetric and Asymmetric groups. The larger proportion of 6 y.o. having Mild hearing loss, and the higher mean thresholds of the 6 y.o. compared with older children, may support these latter alternatives (Table 1). To try to control for the variability of the 6 y.o., we repeated the entire analysis excluding the 6 y.o. group. The results (not shown) were similar to those obtained for the full sample on all the measures in Table 2, indicating the 6 y.o. group did not appreciably skew the outcome.

**TABLE 3.** Auditory perception and cognition were weakly correlated with pure tone threshold (PTA). Data from all children with 4 reliable thresholds and scores on each AP and standardized cognitive test.

| Perception: | BM0 | BM50 | FD | SM | SMN | TR | FR | VCV |
| --- | --- | --- | --- | --- | --- | --- | --- | --- |
| Correlation (r) | 0.148 | 0.199 | 0.105 | 0.056 | 0.125 | 0.034 | 0.087 | 0.188 |
| Significance (p) | <0.001 | <0.001 | <0.001 | 0.035 | <0.001 | 0.201 | <0.001 | <0.001 |
| N | 1427 | 1427 | 1382 | 1422 | 1421 | 1416 | 1414 | 1449 |
| Cognition: | Language | Digits | NVIQ | Words | Pseudo-Words |  |  |  |
| Correlation (r) | -0.108 | -0.103 | -0.079 | -0.106 | -0.108 |  |  |  |
| Significance (p) | <0.001 | <0.001 | 0.002 | <0.001 | <0.001 |  |  |  |
| N | 1457 | 1457 | 1457 | 1443 | 1417 |  |  |  |

### Cognitive Performance

For four of five age-standardized cognitive tests, Normal hearing children performed significantly better than those with Mild hearing loss (Digits, Pseudoword Repetition, Reading Words, Reading Pseudowords; Fig. 5, Table 2). Effect sizes were small to medium (d = 0.19 - 0.53). Children with Minimal hearing loss did not show such a decrement on these tasks, although those with Minimal Symmetric loss were significantly poorer than those with Normal hearing on the language task (Pseudoword Repetition; d = 0.24). Performance of children with Asymmetric and Symmetric loss was statistically equivalent on all measures in both hearing impaired groups. Scores on the NVIQ measures followed these general patterns but did not differ significantly between any hearing categories. None of the attention test scores were affected by hearing status (Table 2).

**Fig. 5:**
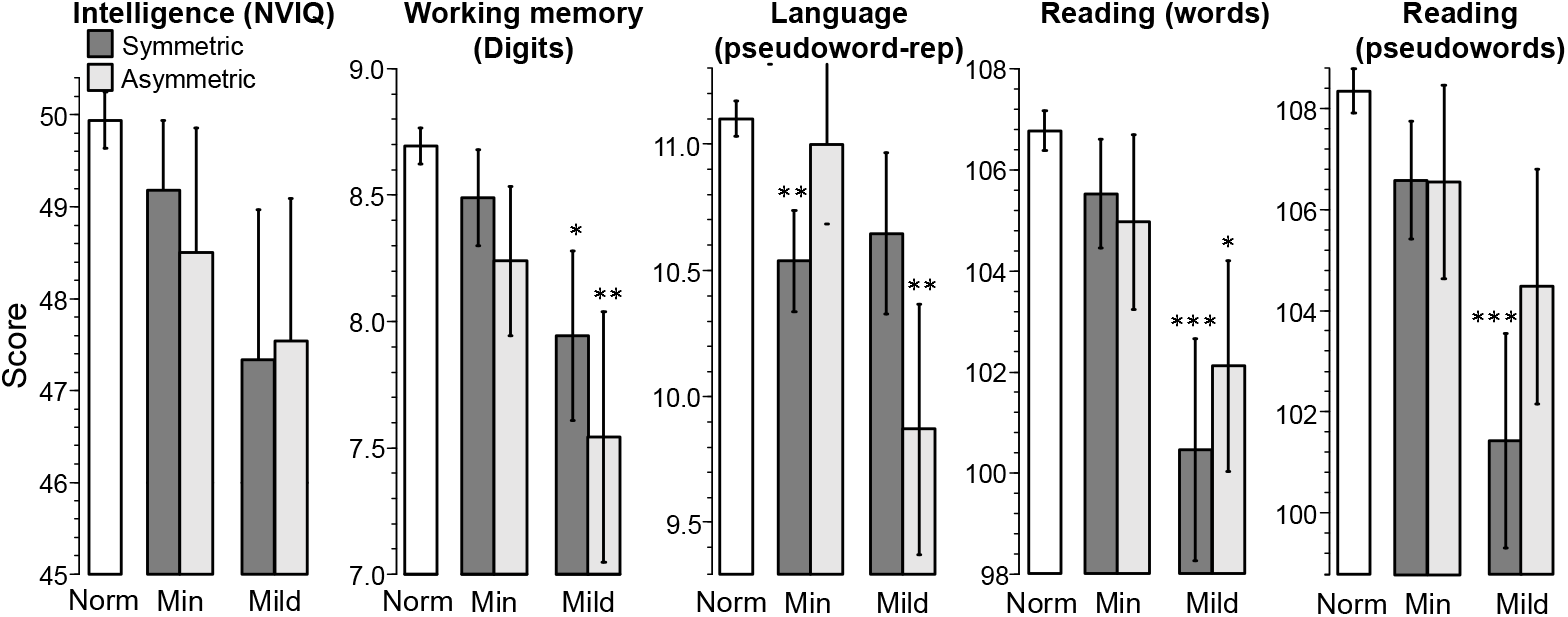
Cognitive performance was reduced in children with hearing loss. Histograms show mean (± s.e.m.) standardized score for each hearing category (details in Fig. 3) and score imputed for missing data. Tests (skill tested) were non-verbal IQ (fluid intelligence), Digit span (working memory), Pseudoword repetition (language), Reading words and pseudowords (literacy). Better performance is shown by higher scores on each test.

Correlations between standardized cognitive test scores and average pure tone threshold across the whole sample (Table 3) were remarkably consistent across tests (|r| = 0.103 to 0.108), with the exception of NVIQ (|r| = 0.079). As for perception, correlations were significant, but weak, and suggestive of multiple other influences on one or both measures.

### Communication and Listening Skills

Communication and listening mean scores of NH children were not significantly higher than those of children with hearing loss (Fig. 6; Table 2). Children with Asymmetric hearing loss performed similarly to those with Symmetric loss. Among Normal hearing children, modest but significant correlations were found between the GCC, 5/8 AP measures (BM50, FD, SMN, VCV, FR; |r| = 0.07–0.19), and all five cognitive measures (r = 0.12– 0.27). For the hearing loss groups, no significant correlations were found between the GCC and the AP tests, except for VCV in symmetric hearing loss (r = 0.30; p = 0.045). However, the much smaller sample sizes of the hearing groups would have contributed to that apparent absence of correlation. Some larger and significant correlations were obtained between the GCC and the cognitive tests (Symmetric: r = 0.24–0.45; Asymmetric: r = 0.03–0.27). Correlations between the CHAPPS, AP and cognitive measures were similar in profile to those of the GCC (e.g. CHAPPS and cognitive measures - Symmetric: r = 0.13–0.47; Asymmetric: r = 0.06–0.39).

**Fig. 6:**
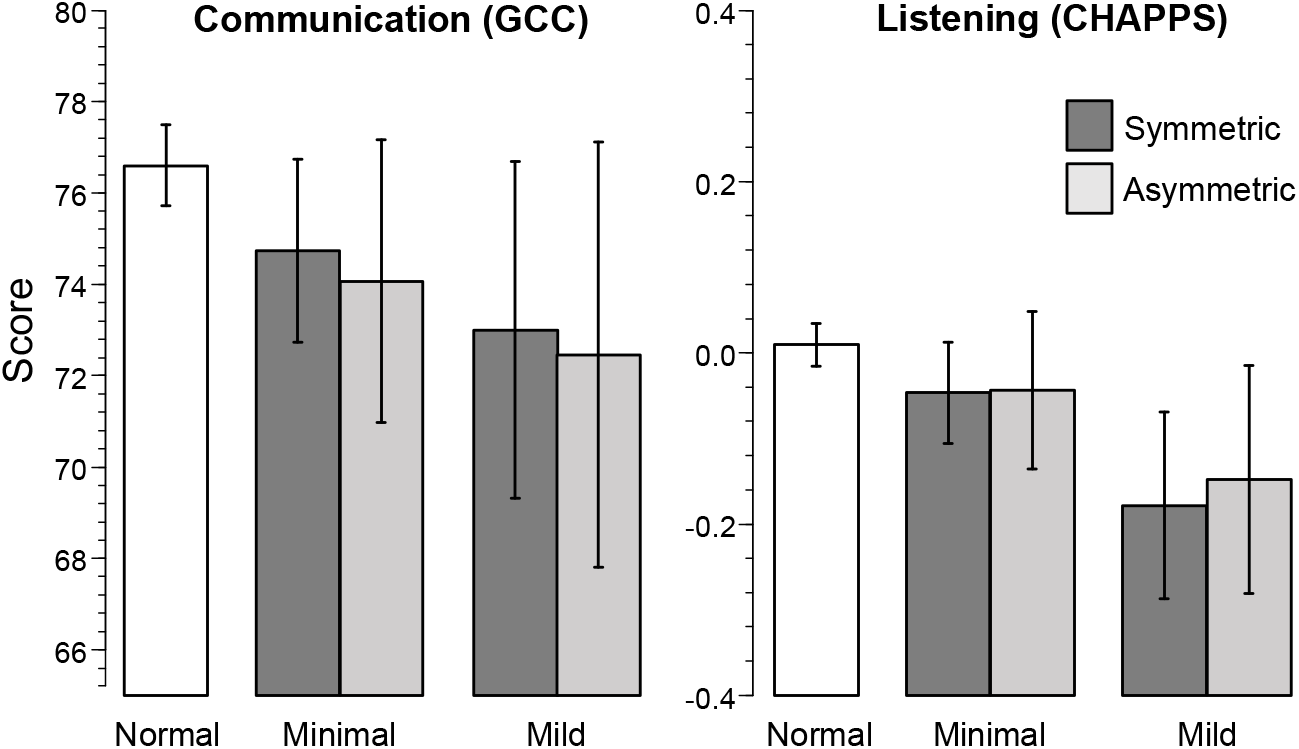
Communication and Listening skills were not statistically reduced in children with hearing loss. Histograms show mean (± s.e.m.) standardized score for each group (details in Figs. 2 and 3). Results were imputed for missing data.

### Additional Clinical Findings

Caregivers reported ear aches/infections, history of otitis media (OM), difficulty hearing, and interactions with hearing health and speech/language professionals (Table 4). We found significant associations between reported frequency of nearly all these events and both Minimal and Mild, and Symmetric and Asymmetric hearing loss (Chi-Square; p < 0.001). Exceptions were visits to speech/language professionals, for whom only modest and mostly non-significant associations with hearing loss were seen. The reported incidence of these issues between children with Minimal and Mild and with Symmetric and Asymmetric hearing loss was mostly similar. However, a smaller proportion of children with Minimal than with Mild loss had reported OM and difficulty hearing faint sounds.

**TABLE 4.** Children with minimal and mild hearing loss had more caregiver-reported hearing problems. A background questionnaire sent to all (n=1622) caregivers at enrolment included the following questions and responses (in percentages). Response rates for these questions ranged from n = 1377 – 1456. Chi-squared tests statistically compared number of children with Symmetric and Asymmetric hearing loss (Minimal and Mild)

| Question: | Has your child had ear aches and/or ear infections? |  |  |  | Has your child seen your doctor (GP) about problems with ears or hearing? |  |  |  |
| --- | --- | --- | --- | --- | --- | --- | --- | --- |
|  | Never | Occasional | Often | Chi-Square p | Never | 1 to 3 | >3 | Chi-Square p |
| Minimal | 40% | 44 | 14 | Min-N: <0.001 | 46% | 33 | 20 | Min-N: <0.001 |
| Mild | 30 | 53 | 16 | Mild-N: <0.001 | 36 | 33 | 21 | Mild-N: <0.001 |
| Symm | 38 | 45 | 15 | Sym-N: <0.001 | 43 | 35 | 22 | Sym-N: <0.001 |
| Asymm | 35 | 51 | 13 | Asym-N: <0.001 | 47 | 33 | 20 | Asy-N: <0.001 |
| Normal | 55 | 45 | 4 | Sym-Asym: 0.60<br>Min-Mild: 0.21 | 62 | 30 | 7 | Sym-Asy: 0.86<br>Min-Mild: 0.64 |

| Question: | Has your child had "glue ear" or chronic otitis media? |  |  |  | Has your child ever visited a hospital because of hearing problems? |  |  |
| --- | --- | --- | --- | --- | --- | --- | --- |
|  | Never | Occasional | Often | p | No | Yes | p |
| Minimal | 81% | 8 | 6 | Min-N: <0.001 | 77% | 23 | Min-N: <0.001 |
| Mild | 66 | 11 | 13 | Mild-N: <0.001 | 68 | 32 | Mild-N: <0.001 |
| Symm | 74 | 10 | 7 | Sym-N: <0.001 | 77 | 23 | Sym-N: <0.001 |
| Asymm | 84 | 5 | 7 | Asym-N: 0.010 | 68 | 32 | Asy-N: <0.001 |
| NH | 91 | 4 | 2 | Sym-Asym: 0.22<br>Min-Mild: 0.02 | 89 | 11 | Sym-Asy: 0.09<br>Min-Mild: 0.10 |

| Question: | Has your child had difficulty hearing faint sounds? |  |  |  | Has your child ever seen a Speech/Language Therapist? |  |  |
| --- | --- | --- | --- | --- | --- | --- | --- |
|  | Never | Occasional | Often | p | No | Yes | p |
| Minimal | 67% | 15 | 11 | Min-N: <0.001 | 84% | 16 | Min-N: 0.03 |
| Mild | 40 | 21 | 17 | Mild-N: <0.001 | 84 | 16 | Mild-N: 0.14 |
| Symm | 62 | 17 | 13 | Sym-N: <0.001 | 85 | 15 | Sym-N: 0.10 |
| Asymm | 65 | 15 | 11 | Asym-N: <0.001 | 82 | 18 | Asym-N: 0.03 |
| NH | 80 | 10 | 3 | Sym-Asym: 0.68<br>Min-Mild: 0.02 | 89 | 11 | Sym-Asy: 0.44<br>Min-Mild: 0.94 |

### Modelling

To investigate in more detail the effect of PTA on outcome, logistic regressions were fitted to AP, cognitive, and caregiver report data for all children (n = 1457; Fig. 7). By several measures, the probability of ‘low performance’ (< 5% of mean Normal hearing performance; see Supplementary information) began to increase above the expected probability (0.05) between 10 - 15 dB HL, where 15 dB HL is the level commonly used to define a ‘minimal’ hearing loss (see Introduction). Cognitive skills (working memory, language and pseudo-word reading) were particularly sensitive to hearing loss, with word reading and NVIQ moderately sensitive. Overall, AP skills were less sensitive to hearing loss, with speech-in-noise the only skill to show moderate sensitivity. Some measures, GCC (Fig. 7), and BM0, BM50, SM, SMN (not shown) had a slight upward trend with increasing level, again from about 10 dB HL, while others (CHAPPS, FD, TR, FR) were relatively unaffected over the range of mean HL (0–25 dB) for which sufficient data were available. The results are generally consistent with Figs. 4 and 5 and demonstrate a range of cognitive and speech-in-noise deficits in children that begin with Minimal hearing loss and become more ‘across-the-board’ in Mild hearing loss.

**Fig. 7:**
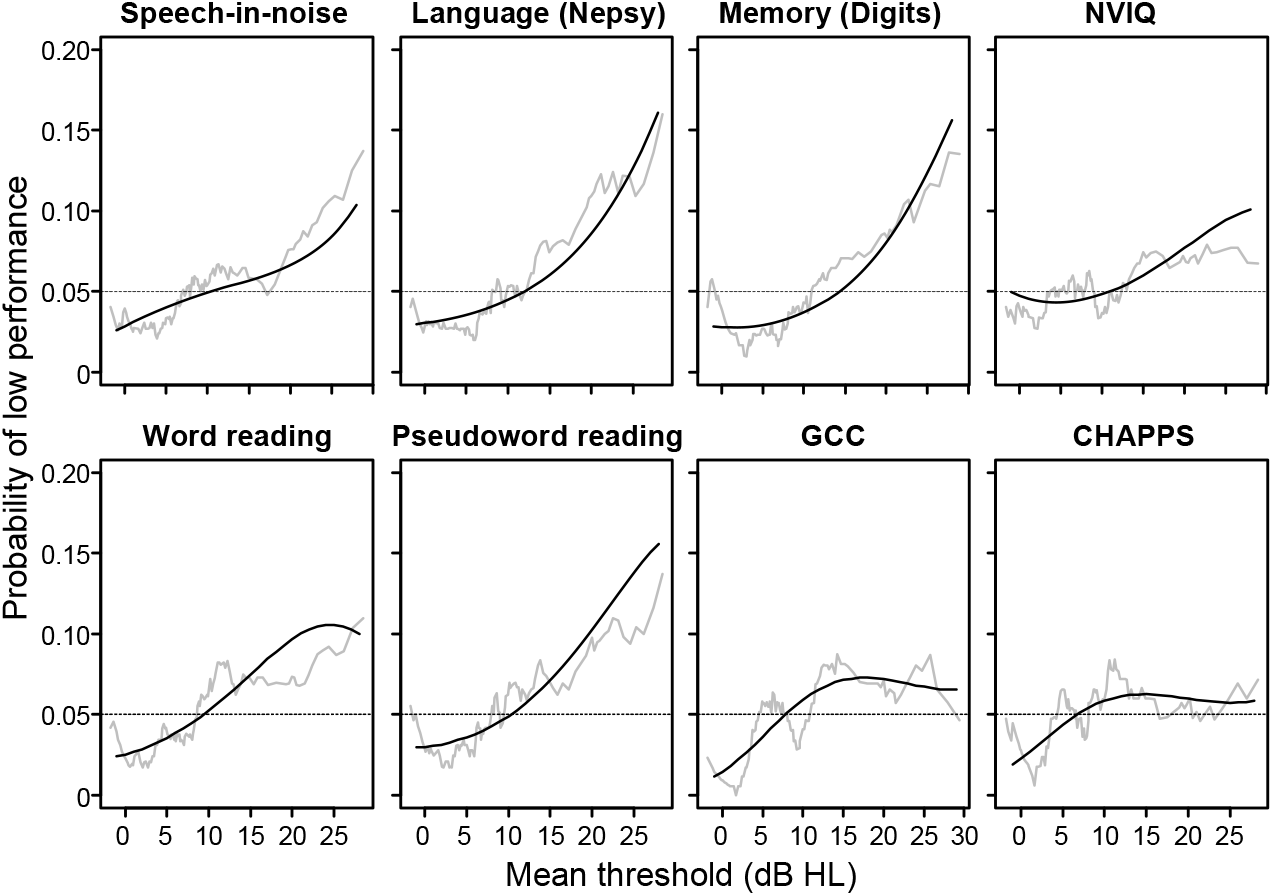
Cognitive and Speech-in-noise performance declined progressively for HL>15dB. Raw data (red lines, moving window average) and fitted data (black lines, cubic logistic regression) indicate the probability of a score for each test that was in the lower 5^th^ percentile of scores on that test in the whole analysed sample (N = 1427; See Supplementary Information for further details).

Addressing possible functional compensation for hearing loss (aim iii), we asked whether listeners with enhanced cognition were less impaired by increasing hearing loss than those with poorer cognition. However, a simple association between AP and cognition among the children with hearing is not evidence for a compensation effect. To establish the presence of such an effect, the benefit of improved cognition should be larger in children with hearing loss than in Normal hearing children. Regression modelling, controlling for age and gender, tested this prediction of deteriorating AP with PTA (HL), but improving AP with NVIQ and Memory (Digits):

> AP~Age+Age^2+Gender+PTA+NVIQ+Memory+NVIQ:PTA+Memory:PTA

Compensation would be indicated by negative interaction regression coefficients, reflecting a strengthening of the relationship between AP and cognition with increasing PTA. However, while confirming the association between AP and cognition, there was no evidence of significant interactions (NVIQ:PTA, Memory:PTA), suggesting a lack of compensation. Further details of these models and associated data are in Supplementary Information.

## DISCUSSION

### Overall findings

Both Minimal (15 - 20 dB HL) and Mild (20 – 40 dB HL) hearing loss in 6 - 11 year old children were associated with poorer than normal auditory perception and cognitive function; the greater the loss, the poorer the outcome. Speech perception in noise and cognitive performance were the most impaired skills. Asymmetric loss produced a similar level of impairment as symmetric loss. For those measures showing poorer than normal performance over the range 0 – 25 dB HL, performance started to deviate from that of children with more sensitive hearing between 10 - 15 dB HL. Differences between Normal and hearing loss groups were, however, generally small, highly overlapping, and not closely correlated between tests. Parental reports of communication and listening skills did not differ significantly with hearing loss over the range examined. Some individual children with hearing loss achieved very good results on all tests, presumably by more effectively using their hearing (compensating). However, children who compensated in this way did not have enhanced cognitive performance relative to children with Normal hearing. In several respects, children with asymmetric hearing loss performed more poorly than Normal hearing children.

Asymmetric loss differed significantly from normal on more individual tests than symmetric hearing loss. Performance patterns suggested that effects of symmetric and asymmetric hearing loss may be underpinned in part by different brain mechanisms. For one of the ‘derived’ hearing measures (frequency resolution), children with asymmetric Mild loss performed significantly more poorly than Normal hearing children, evidence for a ‘bottom-up’ auditory deficit (Moore, 2012; Dillon, 2014, see Methods). It isn’t clear why asymmetric loss produced a disproportionate effect size on this ability. The assumption is that reduced frequency selectivity evidenced by increased filter width derives from the cochlea (Oxenham & Bacon, 2003). It is possible that an imbalance of filter width between the ears interferes with convergent neural pathways onto normally tone-matched targets in the brainstem.

### Type of hearing loss

A potential concern of the methods used here was limited knowledge of type hearing loss. Although 16.8% of this sample had Minimal and 6.0% had Mild hearing loss, some other population studies suggest the prevalence of permanent sensorineural hearing loss of the level (PTA: 15 – 40 dB HL) and across the age range we studied may be lower than this, in the order of 3-5% (Bess et al. 1998). In neonatal hearing screening, most babies failing the initial screen have a temporary conductive hearing loss due to middle ear fluid (Boudewyns et al. 2011). Up to 3-4 years old, OM prevalence remains around 10-30% but, thereafter, a rapid decline occurs, with only about 5% of children showing multiple episodes of OM at 6 y.o (Halliday & Moore, 2010). Nevertheless, it seems likely that conductive hearing loss (CHL) was present in some of the children with hearing loss in the study reported here, and we did find a higher level of OM history and other hearing problems among those children. We did not use bone conduction audiometry, the recommended clinical test to distinguish sensorineural (SNHL) from CHL. However, 5/6 AP tests were signal/masker tests, where a CHL would affect spectrally matched signal and masker levels equally, and all test tones and speech tokens were well above (70 – 90 dB SPL) detection threshold for all children in our sample. All cognitive tests employed visual or supra-threshold auditory stimulation, bypassing audibility effects of CHL. We suggest therefore that all AP and cognitive outcomes would have been similar with or without transient CHL at testing. Following this logic, the likely inclusion of children with CHL in the hearing loss groups may have *reduced* differences found on the AP and cognitive outcomes between the Normal hearing and hearing loss groups, as the observed results would presumably be an underestimate of the difference between a Normal hearing group and hearing loss groups composed exclusively of children with known SNHL. Communication and listening scores were based on caregiver opinion and the relative effects of CHL and SNHL on those opinions cannot be distinguished.

SNHL affects several aspects of AP through loss of cochlear compression, associated with decreasing temporal and frequency selectivity (Oxenham & Bacon, 2003), consistent with the results. However, both CHL and SNHL can produce longer-term consequences. For example, chronic OM in children can produce changes in binaural hearing that persist after the OM has resolved (Hogan & Moore, 2003) and has also been associated with extended high frequency hearing loss (Hunter et al. 1996). It is possible that these effects and others may have influenced speech-in-noise perception or any of the cognitive outcomes.

Another potential influence on the results is that younger children may have elevated tone thresholds due to issues other than hearing loss, including acoustic calibration, normal development of the auditory system, and an inability to respond reliably in the audiometric task (Werner & Gray, 1998), and self-generated noise (Buss et al. 2016). The increased level of mean threshold in the 6 y.o. group relative to the 11 y.o. group at both frequencies would have contributed to the inclusion of more 6 y.o. children in the hearing loss categories and could have been produced by any of the above factors. However, the elevated mean thresholds were rather small, and there appeared to be little if any effect of age-related factors on the ‘outcome’ results, since excluding the 6 y.o. group did not affect the study conclusions. Nevertheless, these considerations do suggest that hearing level should be considered along a continuum, as in Fig. 7 and that, especially in the youngest children, hearing and other immaturity, as well as hearing loss may contribute to poorer performance.

As a compromise between efficiency and complete measurement, only two frequencies in each ear, 1 and 4 kHz, were used to determine hearing loss. This limitation was constrained in part by the test environment, outside a sound booth, that precluded accurate measurement of lower frequency thresholds, and the need to cover the main speech-sensitive range of hearing. If anything, these measures should underestimate the prevalence of hearing loss, but not seriously, we predict, since it seems uncommon for one of these two frequencies not to be involved in hearing loss in children (Pittman & Stelmachowicz, 2003).

### Importance of minimal and mild hearing loss

In this study, almost entirely restricted to children with PTA < 30 dB HL, we found that families did not report statistically higher levels of communication or listening deficits in children with Minimal or Mild hearing loss when responding to questions in the CCC-2 or the CHAPPS. However, they did report a statistically higher proportion of children had difficulty hearing faint sounds, especially in Mild hearing loss. AP and cognitive deficits associated with hearing loss had small to medium effect sizes. Are these effects of practical importance? As outlined above, children with ‘high-end’ mild hearing loss (> ~30 dB HL) perform more poorly academically and socially than their peers and generally do receive timely treatment, at least in many high income countries when detected by neonatal screening (Fitzpatrick et al. 2014; Lieu et al. 2010; Moeller & Tomblin 2015a). A dramatic increase in elementary grade retention rates for children with SNHL (15-40 dB HL; 29 – 47 %) compared with ‘District norms’ (2 – 9 %) was reported by Bess and colleagues (Bess et al. 1998), but their sample straddled both degrees of hearing loss distinguished in the study reported here and their inclusion criteria, in terms of HL and exclusion of children with conductive hearing loss, were more stringent. Here, we found significant and moderate correlations between PTA hearing level across the whole range examined (−13 to 40 dB HL) and parent reported communication and listening skills. These and other correlation data suggested a stronger functional link between hearing loss and communication than between hearing loss and either AP or cognitive performance.

Data for minimal loss, as defined here, are scarce due to low identification other than in population-based studies. It seems reasonable to assume that a PTA < 30 dB HL remains almost entirely undetected and untreated in infancy, despite neonatal hearing screening. Later, pre-school or school entry screening may detect, and act on mild hearing loss (≥ 20 dB HL PTA) but administration of these services appears to be patchy, across UK counties (Bamford et al. 2007) and US states (Gracy et al. 2018). Prevalence also seems to be very low compared with data collected in this study and in several other studies cited here (e.g. Niskar et al. 1998). For example, in a comprehensive national study, (Bamford et al. 2007) cited a large data set showing that, at school entry, only about 1.2% of children had a permanent uni- or bilateral hearing impairment as determined by follow-up assessment. Unfortunately, it appears that a great many, possibly the majority of children with problematic hearing loss around school entry age are not currently detected (c.f. Table 1). More research on minimal and mild hearing loss (PTA = < 30 dB HL) is needed and efforts should be initiated to detect such hearing loss as early in life as possible.

### Management

For children with higher mild hearing loss (PTA = 30-40 dB HL), strong evidence shows the positive effect on communication and education of properly fitted and regularly used hearing aids (McCreery et al. 2015; Moeller & Tomblin 2015a; Tomblin et al. 2015). In light of the previous discussion, hearing aids or other devices are presumably not currently routinely fit to children with hearing loss (PTA) in either ear < 30 dB HL, mainly due to lack of identification of children with those losses. Alternative interventions including counselling on listening skills (e.g. look at a person speaking to you) and preferred environments (e.g. quiet, non-reverberant) are obvious, simple steps (Shield et al. 2010). Advanced communication devices, many based around mobile phones, provide an expanding array of relatively inexpensive and less stigmatic choices that seem particularly suited to adults with minimal or mild hearing loss (Almufarrij et al. 2019). A similar focus by industry on the needs of the paediatric population would be welcome. A simple, low-tech solution would be to introduce classroom-wide amplification using loudspeakers into all classrooms (Dockrell & Shield, 2012). These could serve both to increase signal-to-noise for the teacher’s voice and to draw the attention of all children.

### Further directions

The exclusive or even primary use of conventional audiometry to index hearing impairment is currently being challenged by several emerging discoveries and constructs including unexplained difficulty hearing speech-in-noise (Pienkowski 2017), ‘hidden’ hearing loss (e.g. cochlear synaptopathy; Liberman 2015), ‘extended frequency’ hearing loss (tones > 8 kHz; Monson et al. 2014), and ‘hearing critical tasks’ (Dubno 2018; Soli et al. 2018). The data presented provide another challenge in the form of novel, objective support for a 15 dB HL ‘entry level’ for hearing loss in children. Minimal hearing loss, as stringently restricted here to PTA = 15 – 20 dB HL, is not routinely recognized in clinical practice or used in the literature, although there have been previous calls for a 15 dB lower limit, especially in children (McFadden & Pittman 2008). In this context, it is noteworthy that the skill most affected by Minimal hearing loss in this study was speech perception in noise.

## Acknowledgements

This research was generously supported by the intramural programme of the MRC, the Nottingham University Hospitals National Health Service Trust, The Oticon Foundation and, during analysis and manuscript preparation, NIH Grant R01DC014078, Cincinnati Children’s Hospital Research Foundation, and the NIHR Manchester Biomedical Research Centre. Our gratitude is extended to Alison Riley, senior audiologist, and Mark Edmondson-Jones, statistician, during data gathering and early analysis of the data. Sonia Ratib, research administrative manager, and five research assistants (Karen Baker, Nicola Bergin, Ruth Lewis, Leanne Mattu, Anna Phillips) collected data from the regional test centres over about a one year period. The senior personnel in those centres (Veronica Kennedy, Juan Mora and Kelvin Wakeham) generously provided their facilities and help with the study. IHR technical and support staff provided substantial assistance with the project - we particularly acknowledge the contributions of Tim Folkard, Victor Chilekwa, Dave Bullock and John Chambers. Mark Lutman (University of Southampton) provided software and advice for the automated audiological screen. Lisa Hunter and Dan Sanes engaged DRM in long discussion about the results of this study and strongly encouraged us to publish them. David Moore is supported by the NIHR Manchester Biomedical Research Centre. Finally, we would like to thank all the children, their caregivers and the schools who participated in this study.

## Abbreviations

AP: auditory perception
APD: auditory processing disorder
CCC-2: Children’s Communication Checklist 2
CHAPPS: Children’s Auditory Processing Performance Scale
CHL: conductive hearing loss
BM0: backward masking with 0-millisecond gap
BM50: backward masking with 50-millisecond gap
FD: frequency discrimination
FR: frequency resolution
GCC: general communication composite
HL: hearing level
IHR: Institute of Hearing Research
IMD: index of multiple deprivation
NVIQ: non-verbal intelligence quotient
PTA: pure tone average
SPL: sound pressure level
SM: simultaneous masking
SMN: simultaneous masking with spectral notch
SNHL: sensorineural hearing loss
SRT: speech reception threshold
TR: temporal resolution
VCV: vowel-consonant-vowel speech-in-noise test
y.o.: years old

## Footnotes

1 Two children designated Minimal hearing loss had PTA = 7 dB HL and 7 had PTA = 9 dB HL. The remainder all had PTA ≥ 10 dB HL

